# The IPDGC/GP2 Hackathon - an open science event for training in data science, genomics, and collaboration using Parkinson’s disease data

**DOI:** 10.1101/2022.05.04.490670

**Authors:** Hampton L. Leonard, Ruqaya Murtadha, Alejandro Martinez-Carrasco, Amica Muller-Nedebock, Ana-Luisa Gil-Martinez, Anastasia Illarionova, Anni Moore, Bernabe I. Bustos, Bharati Jadhav, Brook Huxford, Catherine Storm, Clodagh Towns, Dan Vitale, Devina Chetty, Eric Yu, Fatumah Jama, Francis P. Grenn, Gabriela Salazar, Geoffrey Rateau, Hirotaka Iwaki, Inas Elsayed, Isabelle Foote, Zuné Jansen van Rensburg, Jonggeol Jeff Kim, Jie Yuan, Julie Lake, Kajsa Brolin, Konstantin Senkevich, Lesley Wu, Manuela M.X. Tan, María Teresa Periñán, Mary B Makarious, Michael Ta, Nikita Simone Pillay, Oswaldo Lorenzo Betancor, Paula R. Reyes-Pérez, Pilar Alvarez Jerez, Prabhjyot Saini, Rami al-Ouran, Ramiya Sivakumar, Raquel Real, Regina H. Reynolds, Ruifneg Hu, Shameemah Abrahams, Shilpa C. Rao, Tarek Antar, Thiago Peixoto Leal, Vassilena Iankova, William J. Scotton, Yeajin Song, Andrew Singleton, Mike A. Nalls, Sumit Dey, Sara Bandres-Ciga, Cornelis Blauwendraat, Alastair J. Noyce, The International Parkinson Disease Genomics Consortium (IPDGC) and The Global Parkinson’s Genetics Program (GP2)

**Affiliations:** Laboratory of Neurogenetics, National Institute on Aging, National Institutes of Health, Bethesda, MD, USA; Center for Alzheimer’s and Related Dementias, National Institutes of Health, Bethesda, MD, USA; Data Tecnica International LLC, Glen Echo, MD, USA; German Center for Neurodegenerative Diseases (DZNE), Tübingen, Germany; Department of Clinical and Movement Neurosciences, UCL Queen Square Institute of Neurology, University College London, London, UK; Division of Molecular Biology and Human Genetics, Department of Biomedical Sciences, Faculty of Medicine and Health Sciences, Stellenbosch University, Cape Town, South Africa; South African Medical Research Council/Stellenbosch University Genomics of Brain Disorders Research Unit, Stellenbosch University, Cape Town, South Africa; Department of Neurodegenerative Disease, University College London, London, UK; Great Ormond Street Institute of Child Health, Genetics and Genomic Medicine, University College London, London, UK; The Ken & Ruth Davee Department of Neurology and Simpson Querrey Center of Neurogenetics, Feinberg Feinberg School of Medicine, Northwestern University, Chicago, IL 60611, USA; Department of Genetics and Genomic Sciences and Mindich Child Health and Development Institute, Icahn School of Medicine at Mount, Hess Center for Science and Medicine, New York, NY 10029; Preventive Neurology Unit, Wolfson Institute of Population Health, Queen Mary University of London, UK; Department of Human Genetics, McGill University, Montreal, Quebec, Canada; The Neuro (Montreal Neurological Institute-Hospital), McGill University, Montreal, Quebec, Canada; Department of Neuromuscular Diseases, UCL Queen Square Institute of Neurology, University College London, London, UK; INNCOSYS, Col. Morelos Second Section, 50120 Toluca de Lerdo, México; Institut du Cerveau - Institute of Brain and Spine (ICM), Hôpital Pitié, 47 Bd de l’Hôpital, 75013 Paris, France; Faculty of pharmacy, University of Gezira, Wad Medani P.O. Box 20, Sudan; International Parkinson Disease Genomics Consortium (IPDGC)-Africa, University of Gezira, Wad Medani P.O. Box 20, Sudan; Unit for Psychological Medicine, Wolfson Institute of Population Health, Queen Mary University of London, UK; Center for Advanced Parkinson Research, Brigham and Women’s Hospital, Harvard Medical School, Boston, MA 02115, USA; Translational Neurogenetics Unit, Wallenberg Neuroscience Center, Department of Experimental Medical Science, Lund University, Lund, Sweden; Department of Neurology and Neurosurgery, McGill University, Montréal, QC, Canada, Canada; Department of Neurology, Oslo University Hospital, Oslo, Norway; Unidad de Trastornos del Movimiento, Servicio de Neurología y Neurofisiología Clínica, Instituto de Biomedicina de Sevilla, Hospital Universitario Virgen del Rocío/CSIC/Universidad de Sevilla, Seville, Spain; CIBERNED, Madrid, Spain; South African National Bioinformatics Institute (SANBI), South African Medical Research Council Bioinformatics Unit, University of the Western Cape, Bellville, South Africa; Veterans Affairs Puget Sound Health Care System, Seattle, WA, USA; Department of Neurology, University of Washington School of Medicine, Seattle, WA, USA; Laboratorio Internacional de Investigación sobre el Genoma Humano, Universidad Nacional Autónoma de México, Juriquilla, México; Department of Pediatrics, Baylor College of Medicine, Houston, TX, 77030, USA; University of Southern California, Los Angeles, CA 90007; Department of Genomic Medicine, Lerner Research Institute, Cleveland Clinic Foundation, Cleveland, OH 44195, USA; Department of Molecular Medicine, Case Western Reserve University, Cleveland OH 44106, USA; Department of Neurology With Friedrich Baur Institut, University Hospital of Ludwig-Maximilians-Universität München, Munich, Germany

**Keywords:** Hackathon, Parkinson’s disease, genetics, genomics, open science, collaboration, training

## Abstract

**Background:** Open science and collaboration are necessary to facilitate the advancement of Parkinson’s disease (PD) research. Hackathons are collaborative events that bring together people with different skill sets and backgrounds to generate resources and creative solutions to problems. These events can be used as training and networking opportunities.

**Objective:** To coordinate a virtual hackathon to develop novel PD research tools.

**Methods:** 49 early career scientists from 12 countries collaborated in a virtual 3-day hackathon event in May 2021, during which they built tools and pipelines with a focus on PD. Resources were created with the goal of helping scientists accelerate their own research by having access to the necessary code and tools.

**Results:** Each team was allocated one of nine different projects, each with a different goal. These included developing post-genome-wide association studies (GWAS) analysis pipelines, downstream analysis of genetic variation pipelines, and various visualization tools.

**Conclusion:** Hackathons are a valuable approach to inspire creative thinking, supplement training in data science, and foster collaborative scientific relationships, which are foundational practices for early career researchers. The resources generated can be used to accelerate research on the genetics of PD.

## Introduction

An abundance of Parkinson’s disease (PD) data spanning many different modalities (genetic, transcriptomic, proteomic, epigenomic, clinical, and more) has been generated over the past few years. However, many challenges remain in effectively using and integrating this data to produce meaningful and impactful results. A lack of access to good training resources and limited connections to other researchers can delay progress. These problems are even more evident in underrepresented populations, where a lack of resources may hinder researchers and clinicians from performing necessary research on diverse ancestry populations and building local research capacity. To accelerate research, wide adoption of open science practices is critical. Bringing researchers together to share data, code, tools, and pipelines will help reproducibility and create lower entry points to generate the results needed to make necessary progress in PD research.

With the intention of creating pipelines and tools to facilitate PD genetics research, 49 early-career scientists from 12 countries collaborated in a virtual 3-day ‘Hackathon’ event in May 2021. The event combined scientists from two initiatives, the International Parkinson’s Disease Genomics Consortium (IPDGC)^1^ and the first resource project of the Aligning Science Across Parkinson’s (ASAP) initiative, the Global Parkinson’s Genetics Program (GP2)^2^. IPDGC and GP2 exist to drive forward research into the genetic basis of PD, and for both initiatives, training and collaboration are vital aspects of accelerating and diversifying research efforts. Hackathons are a helpful tool that can help promote this mutual effort, providing a creative and engaging outlet to facilitate networking and team building. The first combined ‘GP2/IPDGC Hackathon 2021’ event provided teams with a choice of nine PD genetics and genomics analysis topics. The focus of the GP2/IPDGC Hackathon was not competition, but training, networking, collaboration, and the development of tools that would be useful to the broader research community.

As part of an open science initiative, the Accelerating Medicines Partnership Parkinson’s Disease Program (AMP PD, https://amp-pd.org/)^3^ aims to identify and validate biomarkers by providing researchers access to a large, harmonized dataset that includes clinical, genomic, transcriptomic, and proteomic data. GP2 has recently partnered with AMP PD to make one space where researchers can access multiple PD cohorts and a range of data with a single sign-on. Both AMP PD and GP2 use Terra^4^, a platform for researchers to access data and run analysis tools (Figure 1). The Terra platform allows analysis to be done directly in the cloud, navigating many data sharing and research governance issues that arise from combining data from different geographical regions. Additionally, cloud analysis allows for ease in collaboration, replication of results, and addresses privacy and data use concerns by preventing download of individual level data. Using Terra is necessary for accessing the wealth of AMP PD and GP2 data, so many of the project topics for this Hackathon involved gaining experience and building pipelines on Terra. This allowed hackathon participants to get comfortable using Terra and cloud computing and make available tools to help future researchers use Terra easily and quickly. Whether the teams employed Terra or another platform to create their tools, all were created with the idea to openly share whenever possible to any interested party to accelerate progress.

**Figure 1:**
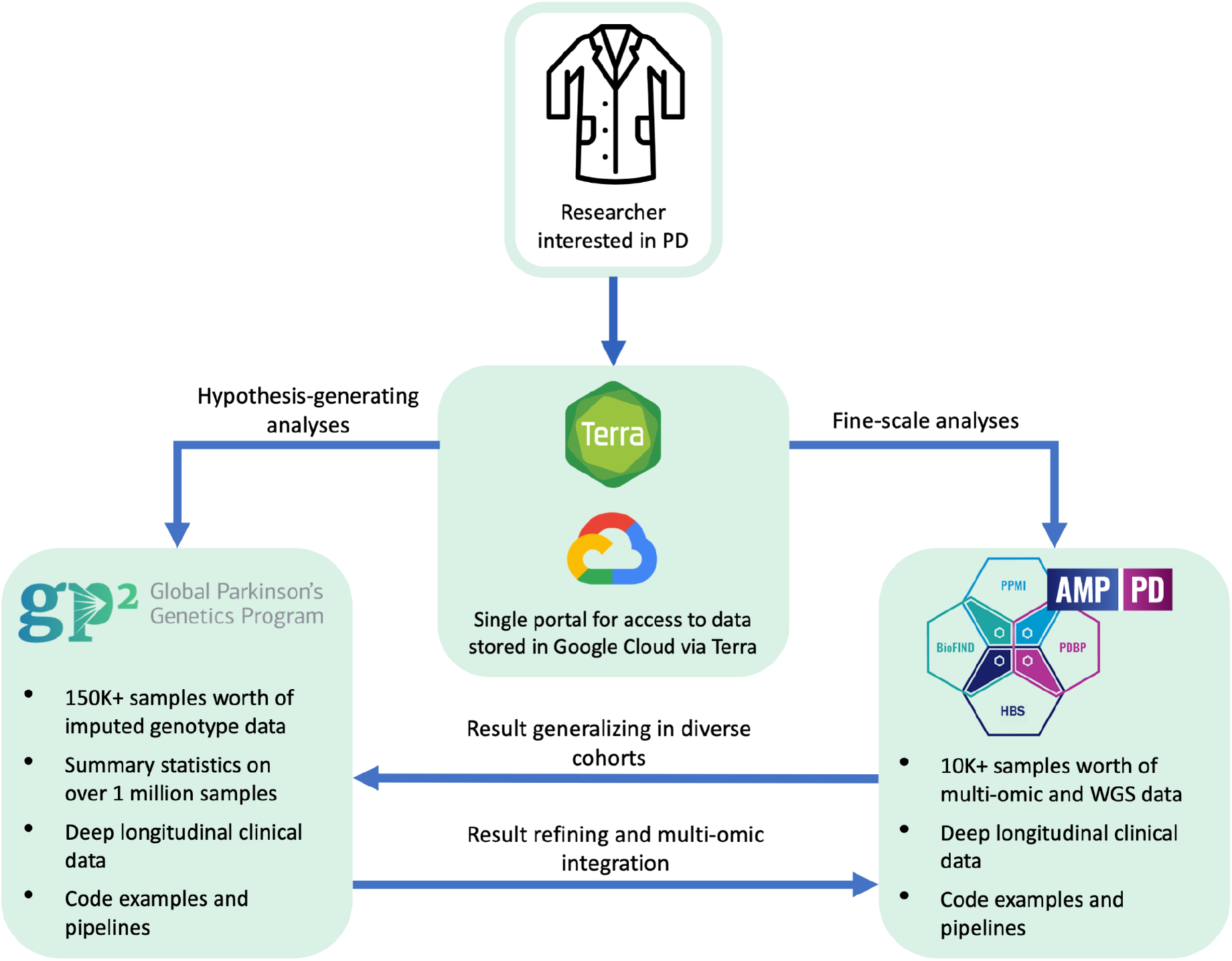
The AMP PD and GP2 collaboration framework. GP2 sample numbers represented in this figure refer to planned numbers, not current. A single data use agreement (DUA) and the Terra cloud platform handle access to both data resources. Each resource addresses different but complementary research questions. GP2 and its wealth of diverse population genotype data are suited for large population-based and hypothesis-generating analyses. In contrast, AMP PD and its multi-omic and whole-genome sequencing data are suited for fine-scale analyses.

## Project summaries (containing both methods and results)

### Overview

To simplify the description of the Hackathon, the nine projects from this event have been grouped into one of three categories; genome-wide association study (GWAS)-level and post-GWAS analyses, downstream analyses of genetic variation, and data visualization. GWAS-level and post GWAS projects were designed to be helpful to researchers looking for ways to follow up their GWAS analyses easily. The projects for this topic included efforts to create general post-GWAS and pathway and cell-type enrichment workflows and examples. Downstream analyses of genetic variation projects aimed at providing examples and pipelines for additional investigation of genetic variation, including colocalization, variant interaction, and network generation and visualization. Finally, the data visualization projects aimed to provide resources that other researchers may use to help further their research or generate hypotheses. These included a GP2 cohort browser, expansion of a PD locus browser, visualization of longitudinal PD clinical outcomes, and visualizing longitudinal and cross-variant effects. A summary of the projects and the topics they belong to and links to available code and applications are found in Table 1.

**Table 1.**
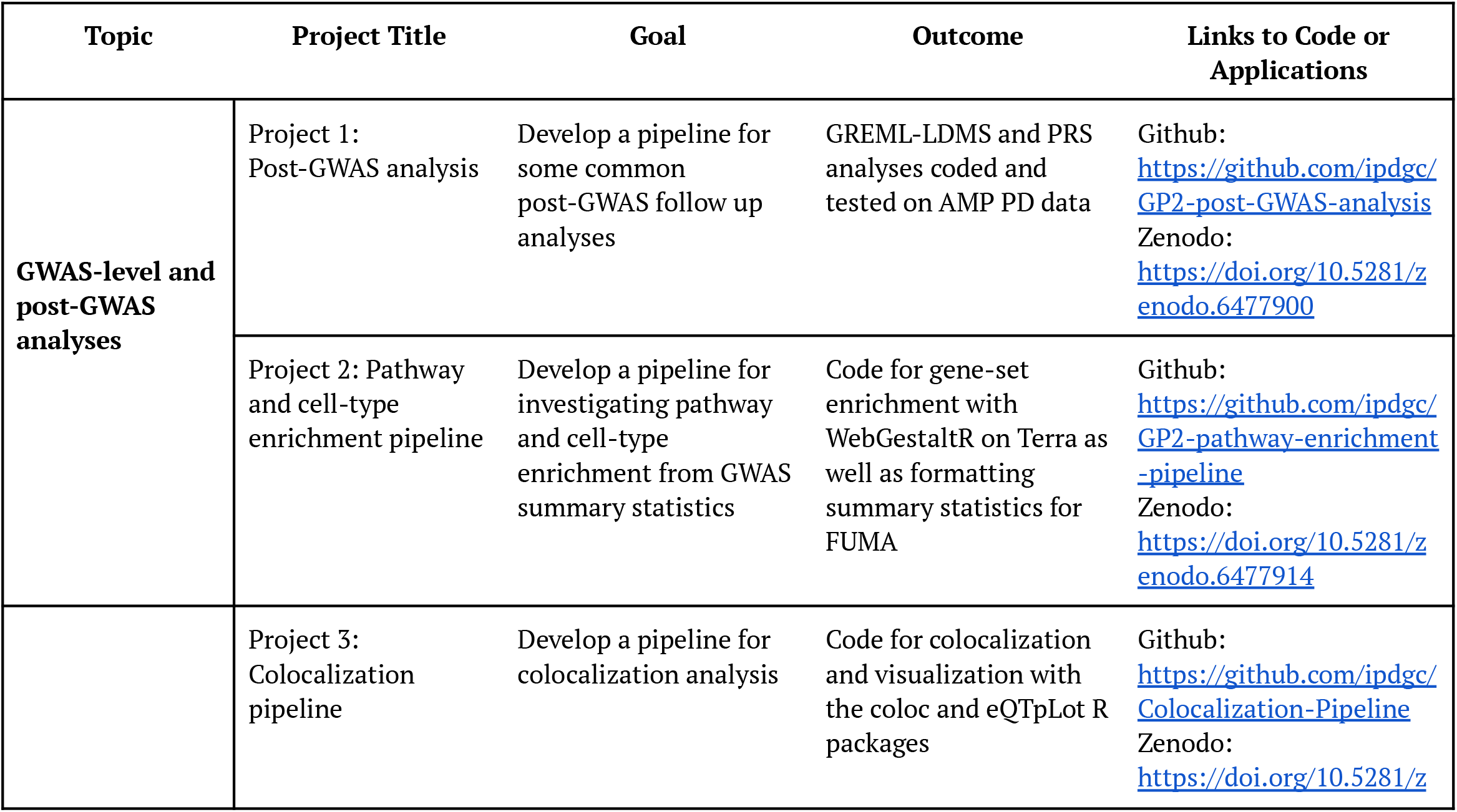
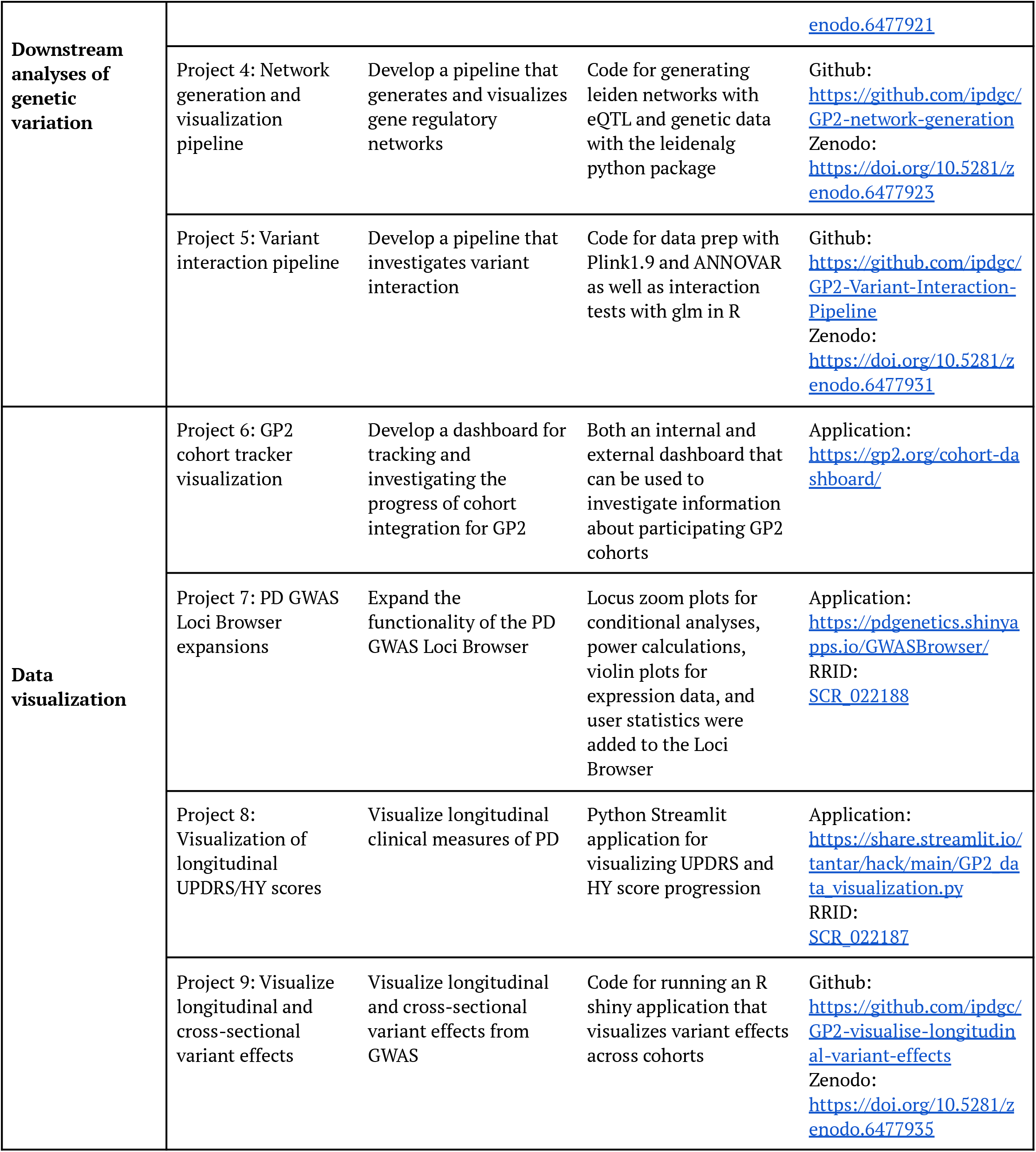
Summary of the goals and outcomes of each of the hackathon projects. Access to the code or applications developed during the hackathon is included in the ‘Links to Code or Applications’ column.

**Table 2:**
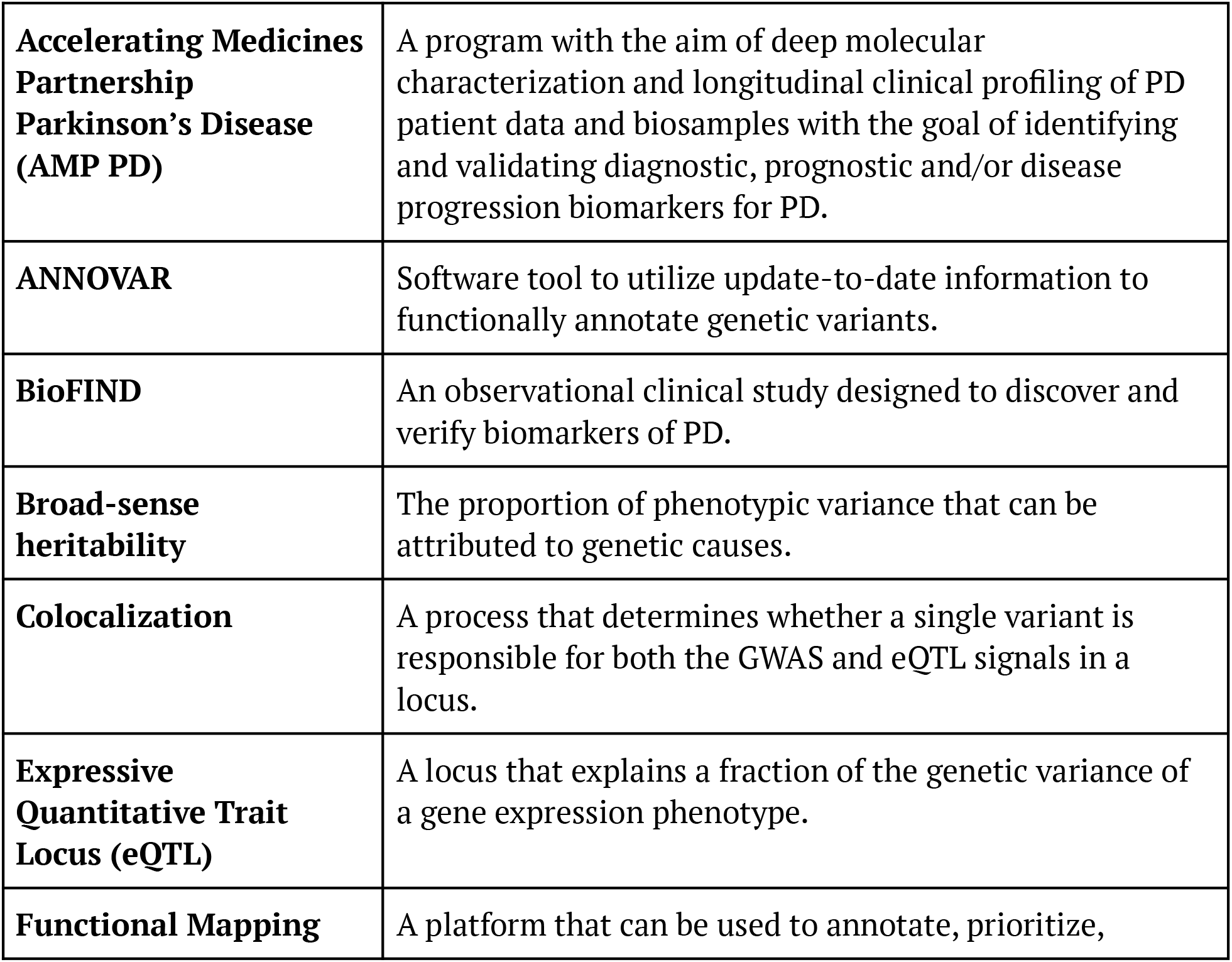
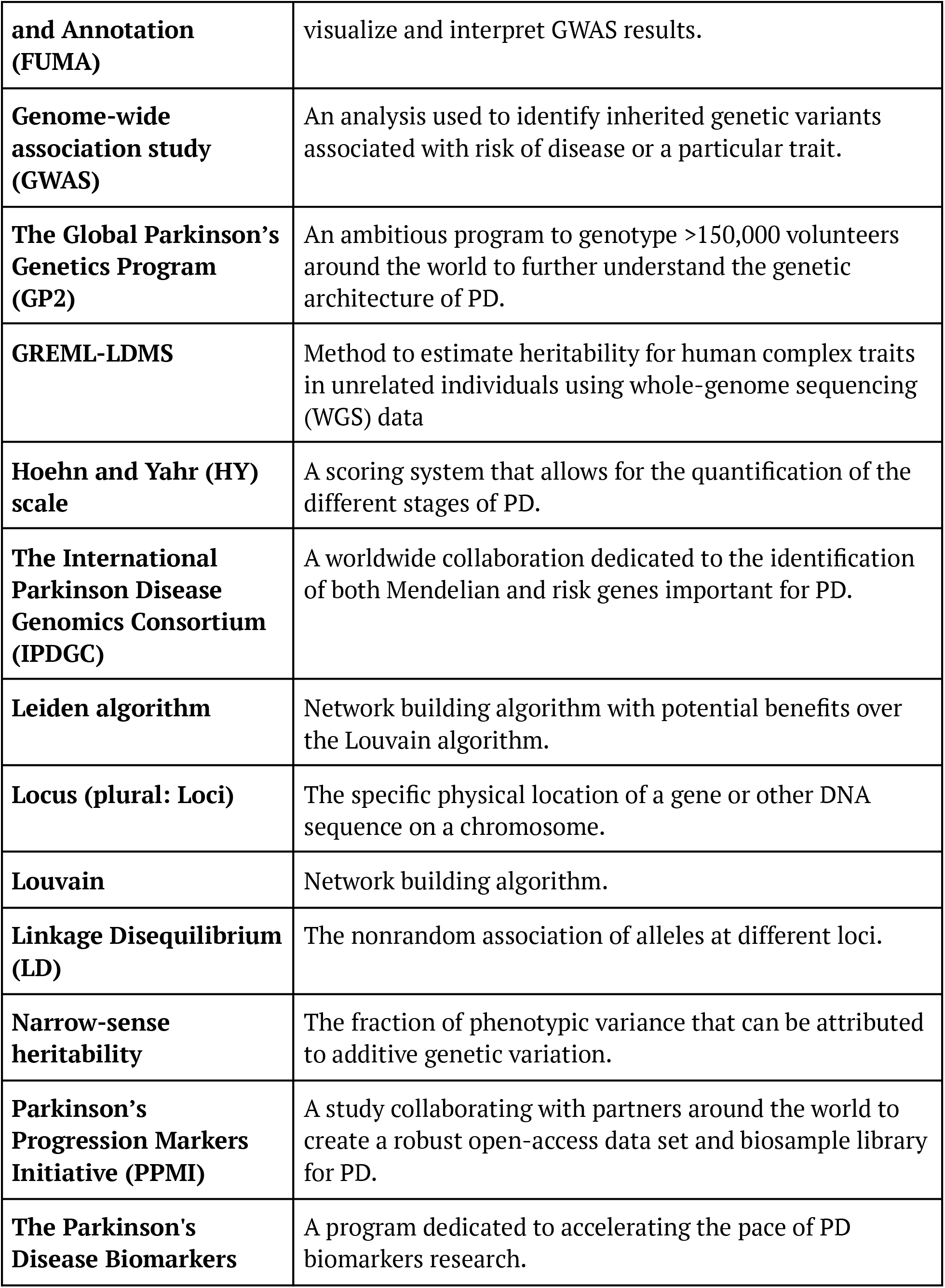
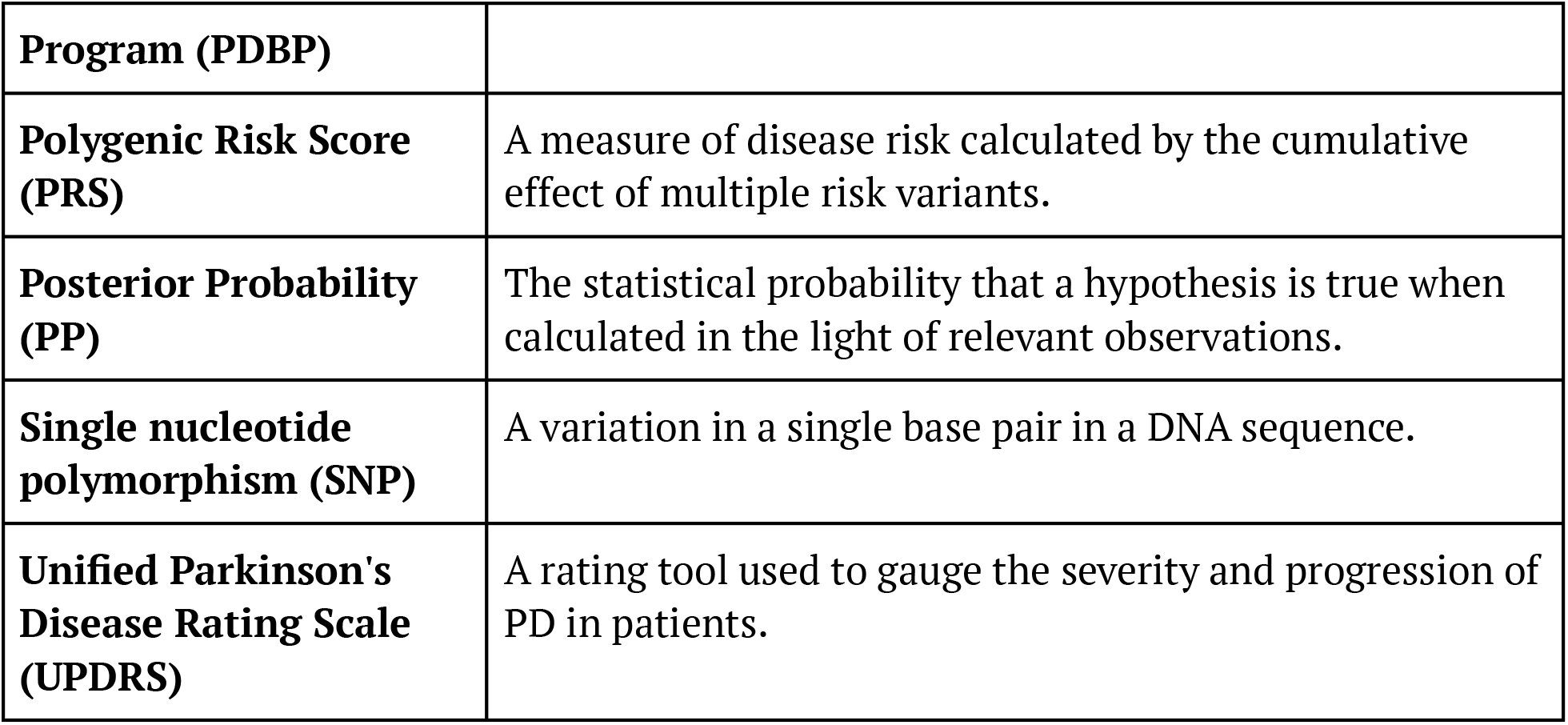
Glossary for unfamiliar terms.

### GWAS-level and post-GWAS analyses

GWAS of PD have nominated 90 independent risk signals in individuals of European ancestry, explaining ∼16–36% of the heritable risk^5^, as well as 2 additional risk signals in Asian populations^6^. Typically, published GWAS are accompanied by various follow-up analyses, but performing these analyses is not always straightforward. Common post-GWAS analyses include heritability estimation and polygenic risk score (PRS) calculation. Heritability analyses estimate the percentage of disease risk accounted for by common genetic variants, and PRS can be used to predict disease risk by aggregating the effects at multiple common risk loci.^7^ Another follow-up approach to help interpret GWAS results is to combine Single Nucleotide Polymorphisms (SNPs) into a group of functionally related genes, such as genes belonging to a single biological pathway or cell type. This is called gene set enrichment analysis and is a widely used approach to examine the cumulative effect of SNPs in a particular biological process and determine whether there are particular pathways, processes, or cell types affected in a disease.

Project 1 (Post-GWAS Analysis): We used AMP PD version 1 release data to develop a Terra-based pipeline for assessing SNP-based heritability, as well as polygenic risk score calculation. Using the GREML-LDMS method^8^ applied to data from the AMP PD version 1 release, we estimated narrow-sense heritability (h^2^) to be roughly 52%, which is much higher than the typical estimate of 22%^5^, and is likely biased due to the recruitment of specific variant carriers present in the AMP PD cohort. AMP PD provides information regarding the recruitment arm. Researchers should investigate this information to determine if specific samples need to be removed from certain analyses in future work. We then used PLINK v1.9^9^ and estimated risk effect sizes from the summary statistics of the latest PD GWAS^5^ to calculate the genetic risk scores of PD *LRRK2* mutation carriers (n = 382), control *LRRK2* mutation carriers (n = 275), and control individuals without PD causing mutations (n = 3435) from the AMP PD version 2.5 dataset. We tested normalized z-scores for association with *LRRK2* carrier disease status. Mean ± standard deviation of unadjusted PRS scores were higher in PD *LRRK2* mutation carriers (−0.0166 ± 0.004), compared to control *LRRK2* mutation carriers (−0.0182 ± 0.004) and controls (−0.0180 ± 0.004), suggesting that PD *LRRK2* mutation carriers share a common polygenic risk profile with idiopathic PD (iPD), contrary to control *LRRK2* mutation carriers (Figure 2A).

**Figure 2:**
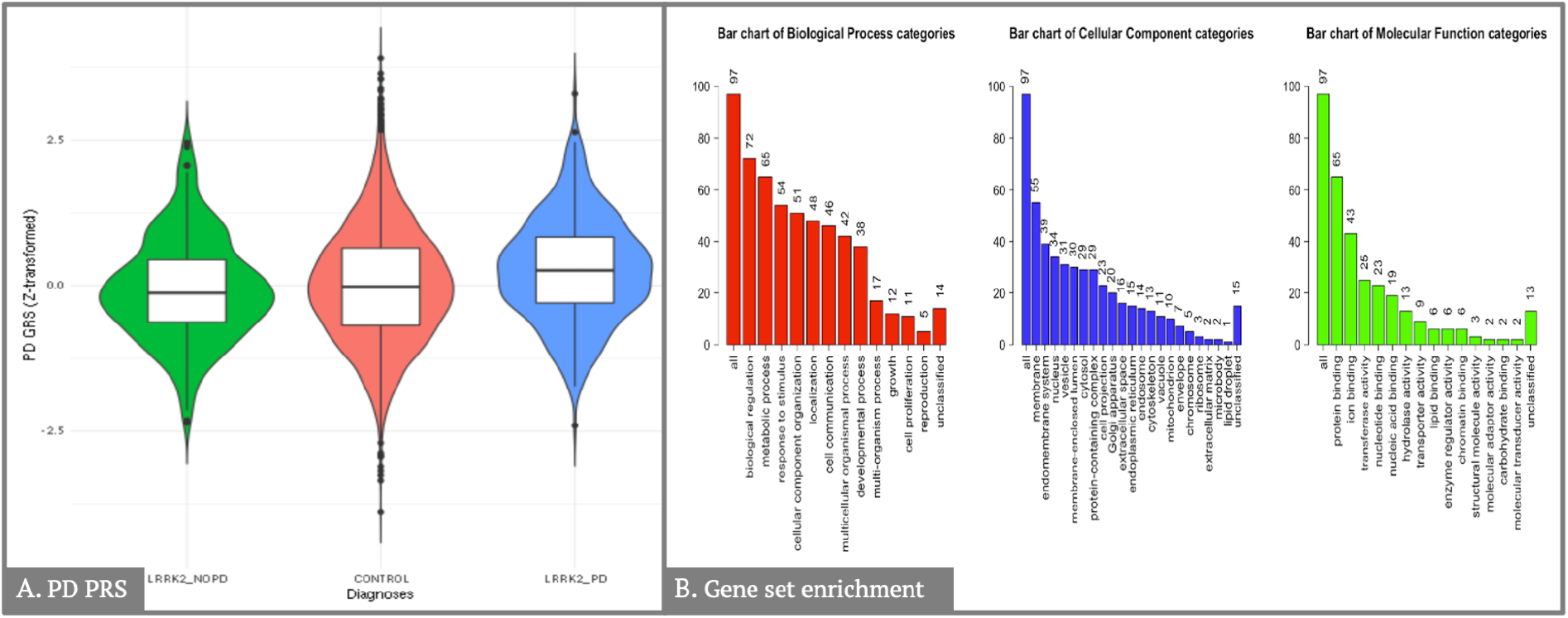
Results from the GWAS-level and post GWAS analyses projects **A**: Violin plots comparing *z*-transformed Parkinson’s disease (PD) genetic risk score distributions in PD-*LRRK2* cases, non-PD-*LRRK2* carriers and controls. PD-*LRRK2* individuals had a higher risk for developing PD compared to control LRRK2 mutation carriers (OR = 1.60, 95% CI = 1.33-1.93, *P* = 1.1×10^−06^). The mean of the unadjusted GRS score was also significantly higher in PD-*LRRK2* cases compared to non-PD-*LRRK2* carriers (*P* = 2.9×10^−06^) and controls (*P* = 5.1×10^−07^) in the pairwise Wilcoxon rank sum test. **B**: Summary of input genes from WebGestalt showing the number of PD genes (from GWAS significant SNPs) which overlap with the annotated genes in the Gene Ontology Slim terms from biological process, cellular component, and molecular function.

Project 2 (Pathway and cell enrichment pipeline): We aimed to create a pipeline to annotate GWAS summary statistics to test the enrichment of biological pathways and cell types. Analyses were performed on the Terra platform using the most recent PD GWAS summary statistics^5^. We created a pipeline to correctly format the GWAS summary statistics for a common annotation tool Functional Mapping and Annotation of Genome-Wide Association Studies (FUMA)^10^, then downloaded the formatted data and uploaded it to FUMA. We also ran the WEB-based GEne SeT AnaLysis Toolkit (WebGestalt) directly on Terra using the WebGestaltR package^11^. We selected the nearest genes to the GWAS significant SNPs (*P* < 5×10^−8^) from Nalls et al. 2019 GWAS summary statistics. Using WebGestaltR, we conducted Overrepresentation Analysis and Gene Set Enrichment Analysis. We identified 97 unique genes from the genome-wide significant hits in the PD GWAS summary statistics. We generated summary data for these PD genes annotated by biological processes, cellular components, and molecular functions (Figure 2B). There was no significant enrichment of any Kyoto Encyclopedia of Genes and Genomes (KEGG) pathway gene sets in the Overrepresentation Analysis (FDR *P* < 0.05).

### Downstream analyses of genetic variation

While GWAS have identified many common variants associated with complex diseases like PD, it is follow-up analyses that have started to decode GWAS results, and more downstream analysis is needed to unravel the implications of observed genetic variation in PD. Three types of analysis were the focus for the downstream analyses of genetic variation topic, including colocalization, variant interaction, and network generation and visualization. Colocalization analysis allows calculation and estimation of the correlation between a GWAS locus and an expression quantitative trait locus (eQTL). Variant interaction, or epistasis, is an interaction of genetic variation at two or more loci to produce a phenotypic outcome that is not predicted by the additive combination of effects attributable to the individual loci^12^. Its importance in humans continues to be a matter of debate^13,14^, but it may explain some of the “missing heritability” underlying complex diseases such as PD^14–16^. In addition to investigating individual variant effects with colocalization and epistasis, visualizing biological networks can help with understanding complex molecular relationships and interactions. In PD research, genetic and gene expression data has been used in community network analysis to nominate pathways and genes for drug target and functional prioritization^17,18^.

Project 3 (Colocalization pipeline): Colocalization analysis takes into account five hypotheses: H0 (no association between the locus and either trait), H1 (locus has an association with first trait only), H2 (locus has an association with second trait only), H3 (locus has an association with both traits but driven by different SNPs which are not in linkage disequilibrium (LD)), H4 (locus has an association with both traits driven by same SNPs). For Project 3: Colocalization pipeline, we considered colocalization analysis with a posterior probability of colocalization in H4 (PPH4) greater than 0.8 to be significant. We utilized the coloc R package^19,20^ and summary statistics from Nalls et al. 2019^5^. We used eQTL data from a cerebellar cortical meta-analysis of four cohorts^21^, publicly available from the AMP-AD Knowledge Portal^22^. As an example for our pipeline, we extracted the region +/- 500 kb around *DYRK1A*, nominated in Nalls et al. 2019, from the GWAS summary statistics and eQTL data. To visualize the results, we employed the eQTpLot R package^23^, which can generate different plots for GWAS and eQTL signal colocalization, as well as the correlation between their p-values and enrichment of eQTLs among variants and LD of loci of interest, allowing efficient and intuitive visualization of gene expression and trait interaction. We used our previously generated results for *DYRK1A* and whole brain eQTL as an example for creating visualizations using this package. (Figure 3A).

**Figure 3:**
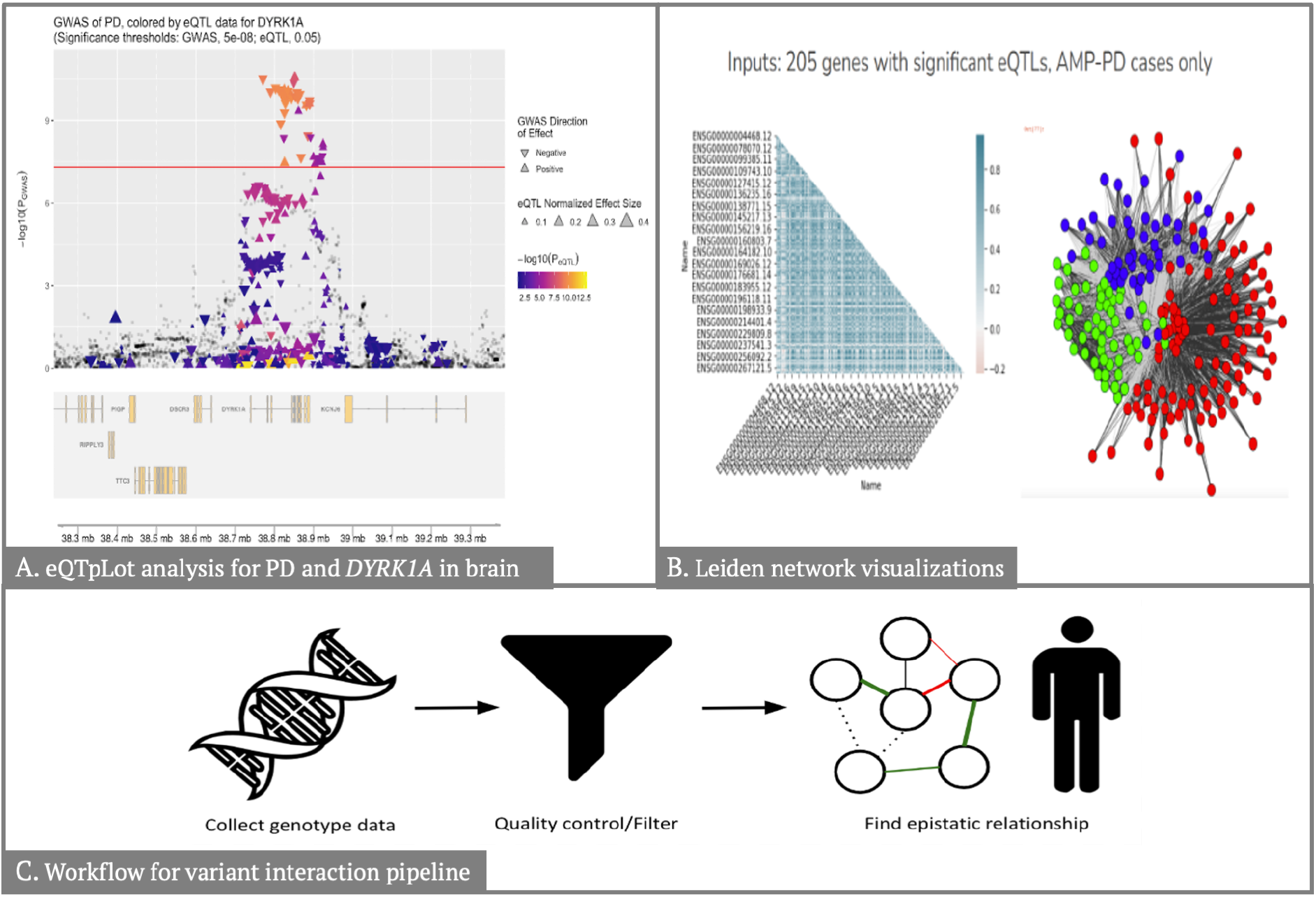
Results from the downstream analysis of genetic variation projects **A:** Displays the locus of interest, in this case, +/- 500 kb from *DYRK1A*, and the horizontal line depicts the GWAS significance threshold of P = 5×10^−8^. Displays the genes in the locus of interest. **B:** Depicts the Leiden gene networks and correlations for significant eQTLs for PD controls and PD cases. **C:** Depicts the general workflow for the variant interaction pipeline.

Project 4 (Network generations and visualization pipeline): We sought to develop a Leiden network and subsequent visualization pipeline for transcriptomic and genomic data to identify and visualize both *a priori* and complex phenotype gene regulatory networks. The Leiden algorithm is one option for community detection of networks and can be faster and return more reliable results than the more well-known Louvain algorithm^24^. We relied on the leidenalg^25^ package in Python to produce weighted and unweighted networks on GWAS summary statistics and then visualized the resulting networks. (Figure 3B). Data used consisted of AMP PD genomic, transcriptomic data, and public eQTL data from the eQTL catalog^26^ and PD summary statistics from the most recent PD GWAS^5^. This project was designed as a proof-of-concept for a pipeline for detecting gene networks and relating them to PD phenotype information via GWAS summary stats.

Project 5 (Variant interaction pipeline): We developed a workflow that can be summarized as follows: 1) We utilized individual-level test data in binary format to perform data harmonization with PLINK v1.9^9^ to ensure that risk allele was consistent for all the variants; 2) We established a minor allele frequency (MAF) threshold > 0.05 to subset variants, keeping only common genetic variation; 3) We annotated variants of interest using ANNOVAR^27^, differentiating between coding and non-coding as well as annotated predicted gene consequence; 4) We carried out interaction analyses in R 3.6 using the glm() function and adjusting for age, gender, and the first five components. (Figure 3C).

### Data visualization

Visualization of clinical and genetic data play an essential role in research. It can be used to inform the progress of initiatives like GP2, help researchers to view data in a meaningful way, and generate and corroborate hypotheses. As GWAS and other analyses nominate more PD risk loci, efforts to decode the role of these variants and how they interact with both longitudinal and cross-sectional phenotypes will be needed. Four projects focused on data visualization, including a GP2 cohort tracker, updates to the IPDGC locus browser, visualization of longitudinal clinical phenotypes, and visualization of longitudinal and cross-sectional variant effects.

Project 6 (GP2 cohort tracker visualization): We designed the GP2 cohort tracker visualization to show essential information about cohorts recruited for GP2 and showcase their diversity, geographical location of enrollment, ancestry representation, and additional relevant metadata. We designed this visualization to inform progress and inspire others to contribute to this initiative. In the form of a one-page dashboard developed with the open-source Python software Streamlit, the visualization includes separate maps for complex and monogenic cohorts. It was critical to include easy-to-use search and discovery aspects built into the dashboard. If a user knows the name of a particular cohort, then they can pull up information for that cohort that populates the rest of the dashboard. The user can also filter by general methods such as cohort size or country. This design is used internally and externally on the GP2 website (https://gp2.org/cohort-dashboard/) to inform those interested in the progress of GP2’s cohort integration (Figure 4A).

**Figure 4:**
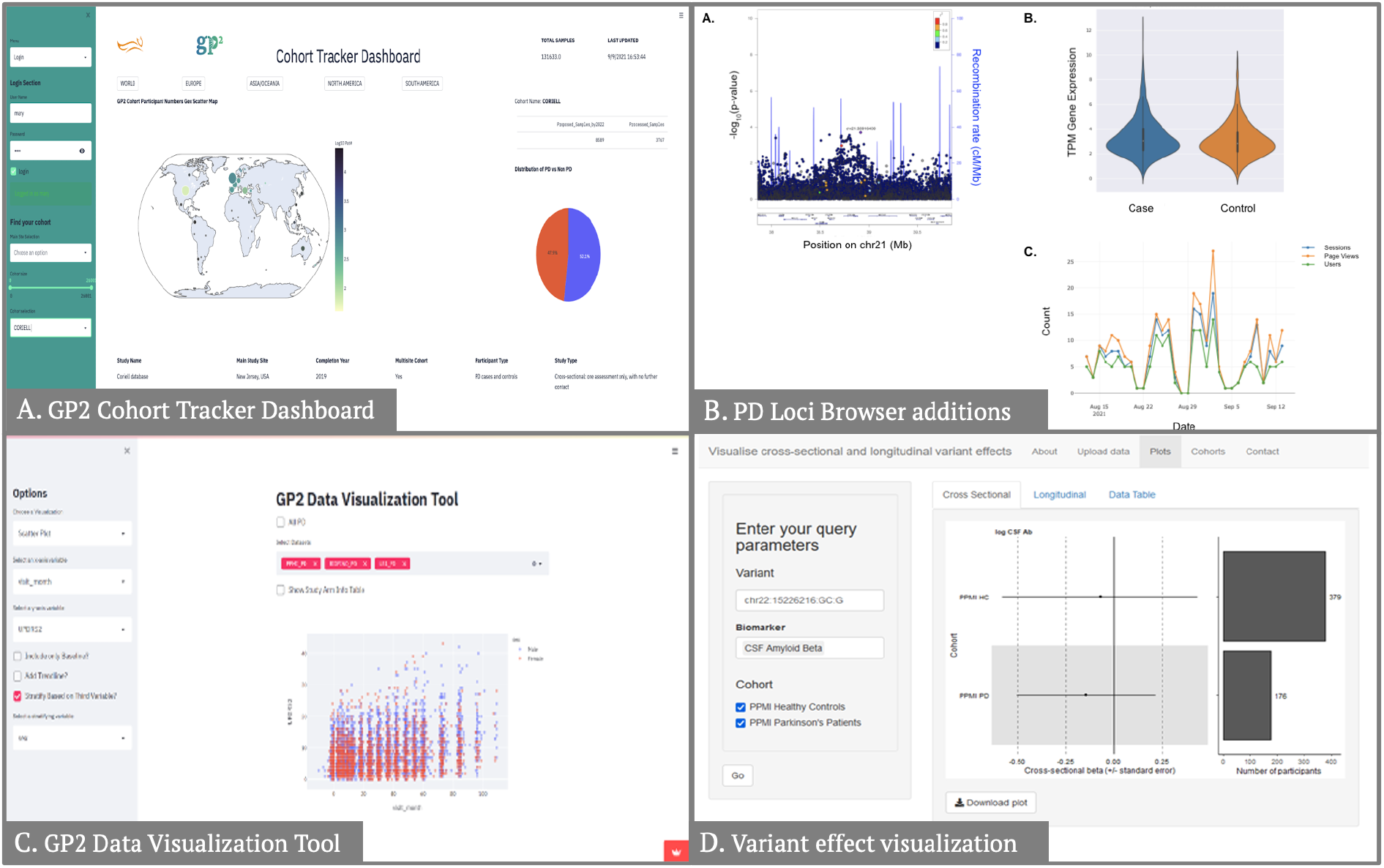
Results from the data visualization projects. **A:** The left banner allows for filtering and specific cohort selection, the map depicts cohort origin, and the right panel depicts PD vs. non-PD distribution. **B:** (A) Locus zoom plot generated using conditional analysis statistics for locus 78 PD risk variant rs2248244. (B) Violin plot of *GBA* expression in AMP PD cases and controls. (C) Example plot of browser user visits over time. **C:** Depicts an example image from the app, in this case, a scatter plot visualization of UPDRS2 scores across visits, color-coded for sex. **D:** On the left, users can input their query parameters, including variant, biomarker(s), and cohort(s) of interest. On the right, a forest plot demonstrates the regression beta for the variant of interest in cross-sectional data. A bar plot demonstrates the number of participants in each cohort. The exact visualization is also available for longitudinal data, with all available data available in a tabular format in the “Data Table” tab.

Project 7 (IPDGC GWAS Loci Browser expansions): To facilitate investigations of nominated risk variants, members of IPDGC have created a PD GWAS locus browser (https://pdgenetics.shinyapps.io/GWASBrowser/) that makes relevant statistics and datasets available to the public^28^. Throughout the Hackathon, our team continued the development of this browser through the addition of new datasets and features. To identify secondary association signals at each locus from the Nalls et al. 2019 study, we performed conditional analysis using the Genome-wide Complex Trait Analysis (GCTA) tool^29,30^. Locus zoom plots were added to display the results of this conditional analysis (Figure 4B)^31^. Power calculations were done for each risk variant from Nalls et al. 2019 to determine if findings were sufficiently powered. To do so, we followed methods used by the Genetic Association Study Power Calculator tool (https://csg.sph.umich.edu/abecasis/gas_power_calculator/), using summary statistics from Nalls et al. 2019, a disease prevalence of 0.01, and a significance level of 0.05 as input. We queried blood gene expression data included in the AMP PD version 2.5 release to measure expression levels in PD cases and controls. We obtained TPM expression at baseline for samples that had case or control status and no PD mutations in whole-genome sequencing data, leaving a total of 1,710 samples. Expression data for each gene was displayed in a violin plot and added to the expression section of the browser (Figure 4B). The literature section of the browser was updated to display a description, PubMed hit count, and word cloud plot for each gene within 1 MB of a PD risk variant. Our last addition to the browser was a display of user statistics. We used the googleAnalyticsR package^32^ to record and visualize the number of visits for the browser and each risk variant within a period specified by the user (Figure 4B).

Project 8 (Visualization of longitudinal UPDRS/HY scores): The Unified Parkinson’s Disease Rating Scale (UPDRS) and Hoehn-Yahr (HY) stage are two of the most common measures of severity of PD. We set out to develop a user-friendly and adaptable app to display a diverse set of visualizations of longitudinal UPDRS/HY scores, based on data from GP2, utilizing the Streamlit library from Python. During the Hackathon, we successfully integrated data from three cohorts: Parkinson’s Progression Markers Initiative (PPMI), Parkinson’s Disease Biomarkers Program (PDBP), and BioFIND^33–35^. We were also able to produce four different visualizations (Figure 4C). First, we created bar graphs to visualize changes in scores from the data over time, and we added the option to include baseline patients only. Second, we created line graphs showing confidence intervals for longitudinal changes in HY and UPDRS scores. Our final visualizations were experimental, but we produced proof of concept visualizations with limited options. We created a Sankey graph visualization that better visualized how participants moved between different subsets of the population over time. Lastly, the fourth visualization is a Kaplan-Meier curve showing time to reach a certain threshold within our progression scores.

Project 9 (Visualize longitudinal and cross-sectional variant effects): We set out with the aim of creating an interactive and user-friendly web application that would allow users to (i) visualize the effect of a genetic variant across multiple cohorts using publicly available GWAS summary statistics and (ii) input their GWAS summary statistics for visualization and meta-analysis with existing data. As test data, we used a small subset of results from a study of amyloid-β levels in cerebrospinal fluid derived from healthy control individuals and individuals with PD from the PPMI dataset^33^. Amyloid-β levels were measured at baseline and in follow-up visits; thus, results were available from both a cross-sectional and longitudinal GWAS. Using the R shiny^36^ framework, we produced a skeleton framework for our web application, with several tabs, including: (i) an “Upload data” tab where users could upload and query their data and (ii) a “Plots” tab where users could query the available test data and visualize it. In the “Plots” tab, we allowed users to query by a variant of interest, with the option to choose which biomarker(s) and cohort(s) they wished to visualize. The variant beta was visualized across cohorts and biomarkers using a forest plot, while the number of participants/observations was visualized using a bar plot (Figure 4D). Tabs were available for cross-sectional and longitudinal plots, with the option to download the plots, and finally, data was also made available in a tabular format.

## Conclusion

In the first IPDGC and GP2 joint Hackathon, 49 early career researchers from across the globe came together virtually to create the tools described here. As the amount of data available for disease research grows and becomes increasingly cloud-centric, public tools like these will help reduce the difficulty and time it takes to visualize, analyze, and understand this data. In order to do effective and prompt research, sharing tools and code for analyses must become the standard.

In addition to creating several helpful pipelines and visualization applications for the PD research community, this Hackathon revealed the need for further documentation and training on cloud computing in the disease research field. Many of the tools created during this event are designed to be used in a cloud setting to assist new researchers in analyzing cloud-based data. However, more resources like these will be needed to ensure cloud resources can be used efficiently.

Hackathons are a valuable tool for prototyping new ideas and tools, but they are also helpful for creating new collaborative networks and working relationships among researchers. This event’s virtual setting allowed many people of different backgrounds and skill sets to work together on creative solutions. Encouraging this kind of creative thinking and creating opportunities for trainees to invest in new and necessary skills is integral to facilitating productive research.

## Acknowledgements and COI

Work was supported in part by Global Parkinson’s Genetics Program (GP2), an Aligning Science Against Parkinson’s (ASAP) initiative implemented by The Michael J. Fox Foundation for Parkinson’s Research (https://gp2.org). Data used in the preparation of this article were obtained from the AMP PD Knowledge Platform. For up-to-date information on the study, see https://www.amp-pd.org. AMP PD – a public-private partnership – is managed by the FNIH and funded by Celgene, GSK, the Michael J. Fox Foundation for Parkinson’s Research, the National Institute of Neurological Disorders and Stroke, Pfizer, Sanofi, and Verily. This research was supported in part by the Intramural Research Program of the NIH, National Institute on Aging (NIA), National Institutes of Health, Department of Health and Human Services; project numbers ZO1 AG000535 and ZO1 AG000949, as well as the National Institute of Neurological Disorders and Stroke.

H.L.L. and M.A.N.’s participation in this project was part of a competitive contract awarded to Data Tecnica International LLC by the National Institutes of Health to support open science research. MAN also currently serves on the scientific advisory board for Clover Therapeutics and is an advisor to Neuron23 Inc.

